# Gesture’s body orientation modulates the N400 during semantic integration of gesture and visual sentence

**DOI:** 10.1101/2020.03.10.985234

**Authors:** Yifei He, Svenja Luell, R. Muralikrishnan, Benjamin Straube, Arne Nagels

## Abstract

Body orientation of gesture entails social-communicative intention, and may thus influence how gestures are perceived and comprehended together with auditory speech during face-to-face communication. To date, despite the emergence of neuroscientific literature on the role of body orientation on hand action perception, limited studies have directly investigated the role of body orientation in the interaction between gesture and language. To address this research question, we carried out an EEG experiment presenting to participants (n=21) videos of frontal and lateral hand gestures of five-seconds (e.g., raising a hand), followed by visually presented sentences that are either congruent or incongruent with the hand gesture (e.g., ‘the mountain is *high*/l*ow…*’). All participants underwent a semantic-probe task, judging whether a target word is related or unrelated to the gesture-speech event. EEG results suggest that, during the perception phase of hand-gestures, while both frontal and lateral gestures elicited power decrease in both the alpha (8-12Hz) and the beta (16-24Hz) bands, lateral gestures elicited reduced power decrease in the beta band when compared with frontal gestures. For sentence comprehension, at the critical word whose meaning is congruent/incongruent with the gesture, frontal gestures elicited an N400 effect for gesture-sentence incongruency. More importantly, this incongruency effect was significantly reduced for lateral gestures. The findings suggest that body orientation plays a crucial role in gesture perception, and that its inferred social-communicative intention influences gesture-sentence semantic integration in an interactive manner.

## 1. Introduction

In every day face-to-face communication, individuals deploy multiple ways to express their communicative intentions. These encompass linguistic expressions in the form of verbal speech, as well as a wide range of non-linguistic cues e.g., gesture, facial expression, eye-gaze, or even body orientation and postures. For example, if an individual wants to keep someone else away, except for verbally expressing *go away*, s/he may additionally produce an accompanying gesture to emphasize this communicative intention. This intention may be even enhanced by a negative facial expression, together with a change of body orientation by turning away from the addressee. Notably, despite the wealth and heterogeneity of these communicative cues, the human brain seems to be remarkably good not only in perceiving, but also in an immediate and efficient integration of all of them. Therefore, an interesting question arises concerning the brain mechanism underlying this integrative process.

Of all these communicative cues, gesture and language are among the most intensively investigated with neuroscientific methods. Literature suggests that human brain is able to effortlessly integrate semantic information from these two input channels. With functional magnetic resonance imaging (fMRI), it is suggested that two regions in the left hemisphere, the left posterior superior temporal sulcus and the left inferior frontal gyrus are crucially involved in the semantic combination and semantic integration between gesture and verbal speech (Dick et al., 2012; Green et al., 2009; He et al., 2015; He et al., 2018b; Holle et al., 2010; Kircher et al., 2009; Straube et al., 2011a; Willems et al., 2009). With electroencephalography (EEG) methods such as event-related potentials (ERPs), it has been shown that human brain is able to rapidly integrate semantic representations from gesture and langauge, as reflected by the N400 component. The N400 has been consistently observed for a variety of experimental manipulations, both for language stimuli that were presented either in auditory or visual form (Fabbri-Destro et al., 2015; Özyürek et al., 2007), and for gesture and speech stimuli that occur simultaneously or consecutively, across adults and children (Habets et al., 2011; Kelly et al., 2004; Sekine et al., 2020; Wu & Coulson, 2005). In general, despite the controversy concerning the functional interpretation of the N400 during semantic processing (Bornkessel-Schlesewsky & Schlesewsky, 2019; Kutas & Federmeier, 2011; Lau et al., 2008), the N400 has been established as a marker for semantic integration between gesture and visual or auditory language (see Özyürek, 2014 for review). In addition, recent studies looking at the oscillatory domain of EEG also showed that both the alpha (8-13Hz) and beta (14-30Hz) bands may support the integration between speech and communicative gesture (Biau et al., 2015; Drijvers et al., 2018a, 2018b; He et al., 2015; He et al., 2018b). Interestingly, these frequency bands were also highly relevant to the perception of action and hand gestures in particular (Avanzini et al., 2012; He et al., 2018a; Järveläinen et al., 2004; Quandt et al., 2012). It has been reported that gestures differing in social-communicative intention (goal-directed vs. non-goal-directed) (Hari et al., 1998; Järveläinen et al., 2004) – as well as the level of simulation (Quandt et al., 2012) – elicited differential level of alpha or beta band power decrease. Additionally, these effects may be even modulated by accompanying auditory speech, as in the case of co-speech gesture (He et al., 2018a).

However, despite the emergence of neuroscientific inquiries into semantic integration between gesture and language, the role of body orientation during this process remains largely unclear. In fact, body orientation (e.g., frontal vs. lateral view) not only presents as physical difference between view-points, but may also indicate differential level of social-communicative intention. The social role of body orientation has been investigated intensively with fMRI studies, the results of which show that it differentially affects either facial emotion or communicative intention (Ciaramidaro et al., 2013; Schilbach et al., 2006) by consistently activating the medial prefrontal cortex (mPFC). Notably, the mPFC is a crucial region within the mentalizing network, which supports the perception of social and emotional features and mentalizing others’ social-communicative intentions (Frith & Frith, 2006; Van Overwalle, 2009). More relevantly, the mPFC is even found to be activated while observing hand gestures that differ in the degree of communicative intentions (Trujillo et al., 2019b).

Nevertheless, body orientation may have a direct impact, physically and socially, upon 1) the perception of hand gestures and 2) the semantic processing during gesture-language interaction. However, both research questions remain elusive with current literature. Research in the domain of action observation has shown that, non-communicative hand actions (reaching, grasping, and etc.) observed from egocentric vs. allocentric view-point elicited greater sensorimotor-related alpha or beta band power decrease (Angelini et al., 2018; Drew et al., 2015). This line of research indicates that body orientation (or view-point) has direct impact on hand action perception. However, none of the existing EEG studies have examined this effect in a communicative context: very often, participants from these studies only see a fraction of the actor/actress (typically one hand), but not the entire body. By contrast, daily communicative gestures are clearly richer than reaching or grasping hand actions in terms of semantic representations and social-communicative intentions. Therefore, it remains an open question whether the perception of communicative gestures will be affected by body orientation (or view point) in a similar way when compared to non-communicative hand actions. These caveats have been addressed by previous studies using fMRI. For example, in Straube et al. (2010), a direct comparison between frontal and lateral co-speech gestures activated the mPFC and other regions in the mentalizing network, thus indicating the potential social-cues conveyed by body orientation (see also Nagels et al., 2015; Redcay et al., 2016; Saggar et al., 2014). However, for these communicative gestures, in the EEG domain, whether body orientation affects sensorimotor alpha or beta oscillations remains unclear.

A more intriguing question concerns the role of body orientation on the semantic integration between gesture and langauge. It is fairly established that gesture and language both share comparable neural underpinnings (Xu et al., 2009), and that the two communicative channels interact with each other (Holler & Levinson, 2019). Neural evidence from both M/EEG and fMRI also shows that gesture modulates language processing, most commonly in a facilitative manner (Biau & Soto-Faraco, 2013; Cuevas et al., 2019; Drijvers et al., 2018b; Zhang et al., 2020). Therefore, given the social nature of communicative gestures, it might be hypothesized that gestures differing in social aspects, as in the case of facing vs. not facing the addressee, may differentially affect the semantic processing of gesture as well as the semantic integration between gesture and language. This hypothesis is being supported by emerging behavioral and neuroscientific evidence. For example, it has been shown that the semantic processing of gestures is facilitated when they are categorized as more communicative (Trujillo et al., 2019a). Moreover, the latter hypothesis is supported by a line of research comparing different gesture styles (Holle & Gunter, 2007; Obermeier et al., 2015): with a disambiguation paradigm, the authors reported that the N400 effect on auditory speech for disambiguating gestures was modulated by adding non-straightforward grooming gestures (Holle & Gunter, 2007). Additionally, the N400 can also be modulated when participants watched an actor who more often produces meaningless grooming gestures, when compared to an actor who always produce straightforward non-grooming gestures. Despite providing initial evidence for social influence on semantic processing of gesture and language, however, these previous studies did not directly compare identical gestures differing in body orientation or view-points.

The purpose of the current study was to therefore 1) find out the electrophysiological markers that differentiate gestures with frontal vs. lateral body orientations during gesture perception, and 2) investigate how body orientation impacts upon semantic integration between gesture and language. To these ends, participants were presented with visual sentences in an RSVP (rapid serial visual presentation) manner, primed by videos of frontal and lateral communicative gestures, and their EEG data were recorded for both gesture perception and sentence processing. Additionally, we manipulated the semantic congruency between gesture and sentence, so as to obtain a reliable measure (N400) of semantic integration between gesture and language. This baseline N400 effect allows us to further investigate how it is modulated by the body orientation of gestures. Based on previous research, we hypothesize that frontal vs. lateral gesture observation will result in differential level of either alpha or beta band power decrease. With regard to the N400 effect that signals gesture-sentence semantic integration, we expect to observe a more pronounced N400 for frontal vs. lateral gestures.

## 2. Methods

### 2.1 Participants

Twenty-four participants (15 female, mean age = 24.95, range 19-35) participated in this experiment and were monetarily compensated for participation. All participants were right-handed as assessed by a questionnaire on handedness (Oldfield, 1971). They were all native German speakers. None of the participants reported any hearing or visual deficits. Exclusion criteria were history of relevant medical or psychiatric illnesses. All participants gave written informed consent prior to taking part in the experiment. The data from three participants were discarded because of excessive movement artifacts during recording.

### 2.2 Materials and procedure

A sample of an experimental trial is illustrated in **Figure1C**. A trial started with a fixation mark of 1000ms, followed by a video of an actor performing a hand gesture with two different body orientations (frontal vs. lateral, **Figure1A**). The frontal and lateral videos were adopted from a line of fMRI studies investigating social aspects of gesture processing (Nagels et al., 2015; Straube et al., 2011b; Straube et al., 2010). They were recorded by two video cameras simultaneously when the actor was gesturing. As a result, only the viewpoint differed between the two conditions. All videos lasted 5000ms and did not contain any auditory information. For all videos, different from previous fMRI studies, we blurred the face area of the actor so that potential differences in eye gaze between frontal and lateral videos were not available to the participants. We identified gesture onsets and offsets for all videos: the mean gesture onset time was about 1.10s (SD = 0.29s), and mean offset was about 3.38s (SD = 0.46s). The videos were followed by sentences presented in an RSVP (rapid-serial-visual-presentation) manner, with 300ms per word and 100ms inter stimulus interval (ISI). Sentences were formulated in two experimental conditions, such that they were either clearly congruent or incongruent with the prime video. All sentences were of the same structure as in **Figure 1B**: *Dass der Fisch <u>groß/klein</u> ist, findet Lara* (The fish is <u>big/small</u>, thinks Lara). Therefore, the congruency between the sentence and the prime video becomes only apparent at the critical adjective, and that the critical words do not occur at the sentence-final position (Stowe et al., 2018). We controlled word length and frequency (Wortschatz Leipzig) at the position of the critical words between both congruency conditions. The length of the critical words in the congruent condition was marginally shorter than that of the incongruent condition (6.87 vs. 7.81, t = −1.86, p = 0.065). The frequency of critical words was not different between the two conditions (12.44 vs. 12.53, t = −0.19, p = 0.85). One second after the presentation of the RSVP sentence, a semantic probe word (related or unrelated, 50% each) appeared for a maximum of 4000ms, prompting the participants to judge, as fast as possible, whether the word is semantically related or unrelated to the previous event (irrespective of gesture or sentence). Altogether, we used 32 videos (seventeen iconic gestures and fifteen emblematic gestures) differing in body orientation. For each video, two sets of (congruent/incongruent) sentences were created. Therefore, the 256 (32-videos x 2-orientation x 2-congruency x 2-sets) experimental items were split into two lists of 128 items each. Each participant was presented with only one pseudo-randomized list consisting of 128 experimental items together with an additional 128 filler items of no interests. The fillers items were formed by a combination of 32 different frontal/lateral videos followed by congruent/incongruent RSVP sentence with varying sentence structure. For each participant, an experimental session lasted for around an hour, distributed in eight blocks.

**Figure 1.**
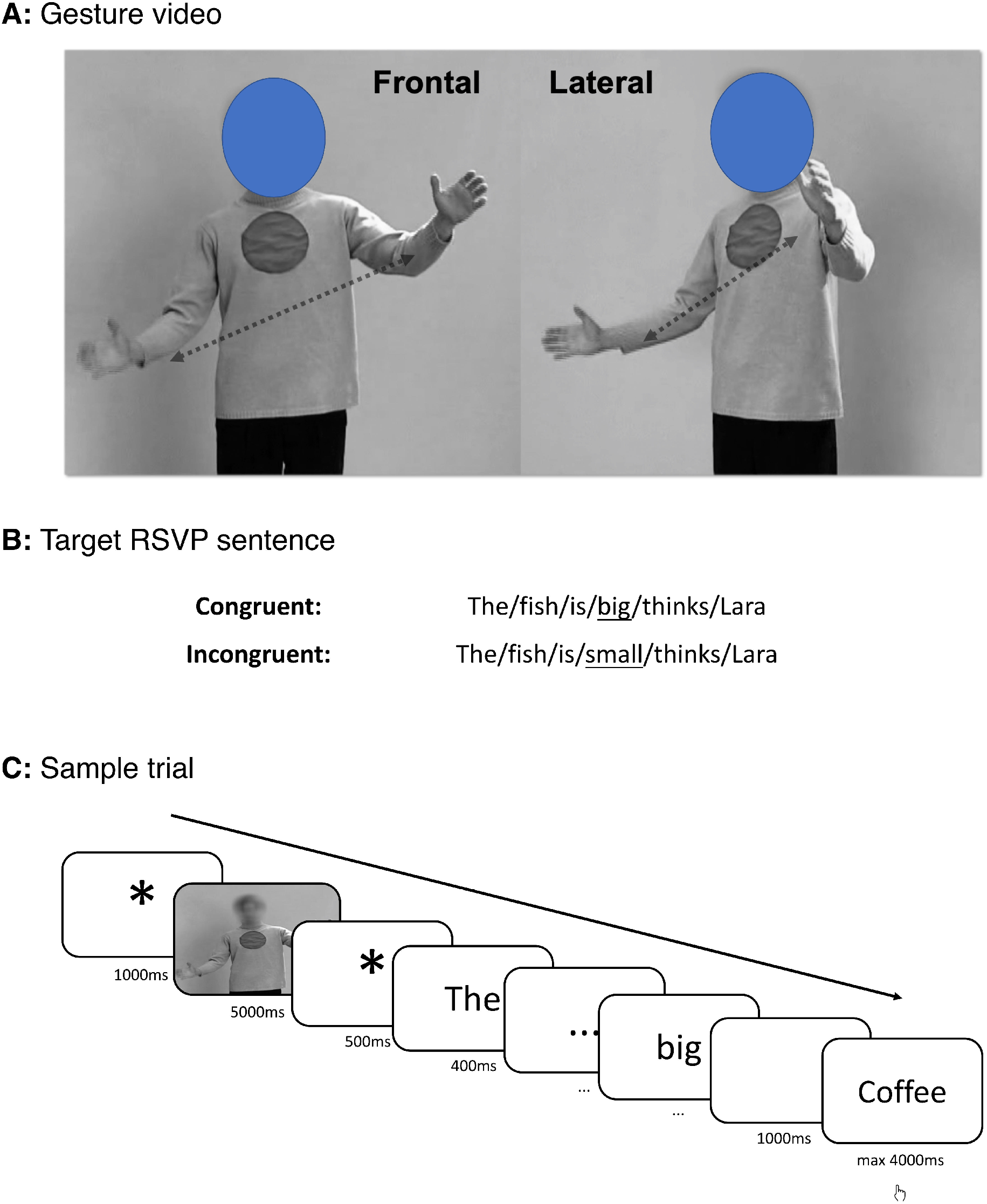
Illustration of the experimental paradigm. Panel A: each communicative gesture was recorded by two cameras and videos were presented with frontal (F) and lateral (L) body orientations. Panel B: congruent and incongruent sentences of an experimental item, the critical word is underlined. Panel C: A sample of an exemplary trial. Each word was presented for 300ms + 100ms ISI. Note that the face of the actor was covered with blue circle under requirement from bioRxiv. Instead, participants watched pixelated face of the actor such that both eye-gaze and mouth movements are unavailable.

### 2.3 EEG acquisition

EEG was collected from 64 Ag/AgCl passive electrodes attached to the an elastic cap (EasyCap GmbH, Herrsching, Germany) according to the international 10-10 System. The reference electrode was located at the FCz and the ground electrode was located at the forehead in AFz. All input impedances were kept below 10 kΩ. Additionally, the vertical electrooculogram (VEOG) was recorded from one electrode located underneath the left eye. Two ‘Brain Amp’ (Brain Products, Garching, Germany) amplifiers were used to sample data at 500 Hz with a resolution of 0.1 μV. Trigger signals from stimulus and participants responses were presented with Presentation Software (Neurobehavioral Systems, Berkeley, CA, USA), and acquired together with the EEG using Brain Vision Recorder (Brain Products GmbH, Garching, Germany).

### 2.4 EEG preprocessing and analyses

All EEG preprocessing and analyses were carried out using the Brain Vision Analyzer 2.1 (Brain Products GmbH, Munich, Germany) and the Fieldtrip toolbox for EEG/MEG analysis (Oostenveld et al., 2011). We applied two processing pipelines at the video onset (comparing time-frequency representation (TFR) between frontal and lateral videos) and at the onset of the critical words in the target sentence (comparing ERPs for the four experimental conditions).

#### 2.4.1 EEG Preprocessing

For the TFR analyses at video onset, raw continuous EEG were firstly high-pass filtered at 0.1 Hz and low-pass filtered at 125 Hz, and then re-referenced to the average of all electrodes. EOG and muscle artifacts were manually identified and rejected via an infomax independent component analysis. The raw EEG was then segmented to segments around - 1 to 5.5 seconds around the onset of video. After segmentation, we firstly applied automatic muscle artifact rejection based on the amplitude distribution across trials and channels (as implemented in Fieldtrip toolbox). Cutoffs for these artifacts were set at z = 6. Afterwards, additional trials were rejected manually. The average rejection rate for both frontal and lateral conditions was 13.61% (SD = 9.48%), with no significant difference between the two conditions.

For ERP comparison time-locked to the critical word onset, we applied a band pass filter between 0.1 to 40 Hz to the raw EEG. The re-referencing and artifacts rejection procedure (for both continuous and segmented data) was analogous to the TFR analyses for video onset, except for the fact that we segmented a shorter time window between −0.2s and 1s based on critical word onset. Data preprocessing resulted in an average rejection rate of 9.63% (SD = 7.62%) for all four conditions, with no significant main effects or interaction between the two experimental manipulations.

#### 2.4.2 TFR and source analyses

As we are primarily interested in the alpha and beta frequency bands for a direct comparison between frontal and lateral videos, for TFR computation, we applied a sliding window Hanning taper approach (5 cycles per window), in frequency steps of 1 Hz in the range of 2-30Hz, and time steps of 0.20 seconds. All TFRs were interpreted based on baseline corrected (−0.5 to −0.25 seconds) decibel change (dB). After time-frequency decomposition, we applied a cluster-based random permutation test (Maris & Oostenveld, 2007) with 1000 permutations to test the difference in both the alpha (8-12Hz) and beta bands (16-24Hz) between frontal and lateral conditions. The test was conducted in the time-electrode space, with each electrode having on average 5.8 neighboring electrodes according to a template layout. A cluster in the permutation test contained at least three neighboring electrodes.

For source localization of the beta band effect (see results), we conducted a frequency-domain adaptive spatial filtering imaging of coherent sources (DICS) algorithm (Gross et al., 2001), as implemented in the Fieldtrip toolbox. Source analysis was performed for the time-frequency windows in the beta band (20Hz with 4Hz padding) in which significant results were obtained on the scalp level. Participants’ electrode positions (defined with a templated electrode layout) were warped to the cortical mesh of a standard boundary element head model (BEM). The forward model was computed on an 8mm grid of source positions covering the whole brain compartment of the BEM. For source analysis, common space filters were constructed using the leadfield of each grid point and the cross-spectral density matrix (CSD). The CSD matrix was computed based on an additional time-frequency analyses for data segments of interests plus their respective baseline. The source activity volumes of interests were firstly corrected against the baseline period, and were then compared by means of paired t-tests between both experimental conditions. This procedure resulted in a source-level t-statistics of alpha power change for each voxel in the volume grid, in which we thresholded at a level of p<0.05, uncorrected. For identification of anatomical labels, we used the AAL atlas (Tzourio-Mazoyer et al., 2002).

#### 2.4.3 ERP analyses

In order to reveal the N400 difference at the critical word, for each segment, we applied a baseline-correction based on the −0.2 to 0 seconds time window. Segments within a condition were then averaged within each participant across trials in the first step. Secondly, grand averaged ERPs were averaged across all participants. For ERPs, we used cluster-based permutation test (1000 permutations) for statistical comparison in the electrode space for amplitudes averaged within the N400 (300-500ms) time window. Neighborhood parameters were set comparable to the TFR analysis. As we were interested in the interaction between body orientation and sentence-gesture congruency, we firstly compared congruency effects within each body orientation conditions, and then compared the incongruent>congruent difference between frontal and lateral conditions to test for statistical significance of interaction.

## 3. Results

### 3.1 Behavioral results

The reaction times and accuracy for the semantic probe task is reported in **Figure 2**. For reaction times, repeated-measures ANOVA showed a significant main effect of body orientation (F_(1, 20)_ = 11.32, p = 0.003). For accuracy, we observed no main effects (F_(1, 20)max_ = 2.71, p = 0.12) but a significant interaction between body orientation and congruency (F_(1, 20)_ = 6.15, p = 0.03). However, pairwise t-tests showed that congruency effects in both frontal and lateral gesture conditions were not significant (|t|_max_ = 1.70, p = 0.10).

**Figure 2.**
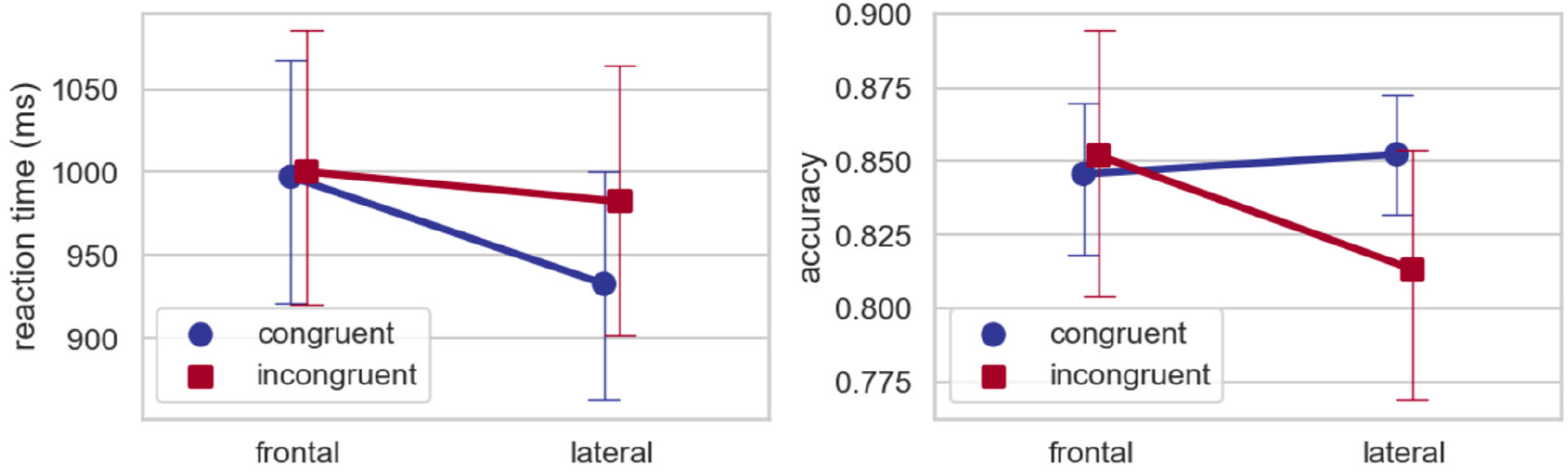
Reaction times (ms) and accuracy for the semantic probe task.

### 3.2 Video onset: beta band oscillations

We directly compared time-frequency representations between frontal and lateral gestures (**Figure 3**). For both conditions, although we observed alpha and beta power decrease relative to the baseline, only the beta band showed significant difference between the frontal vs. lateral conditions (p_cluster_ = 0.005). This effect has a right scalp distribution. The source of this beta band effect was source localized to the right middle frontal cortex (MNI coordinates: x = 34, y = 53, z = 12, |t|_max_ = 2.78).

**Figure 3.**
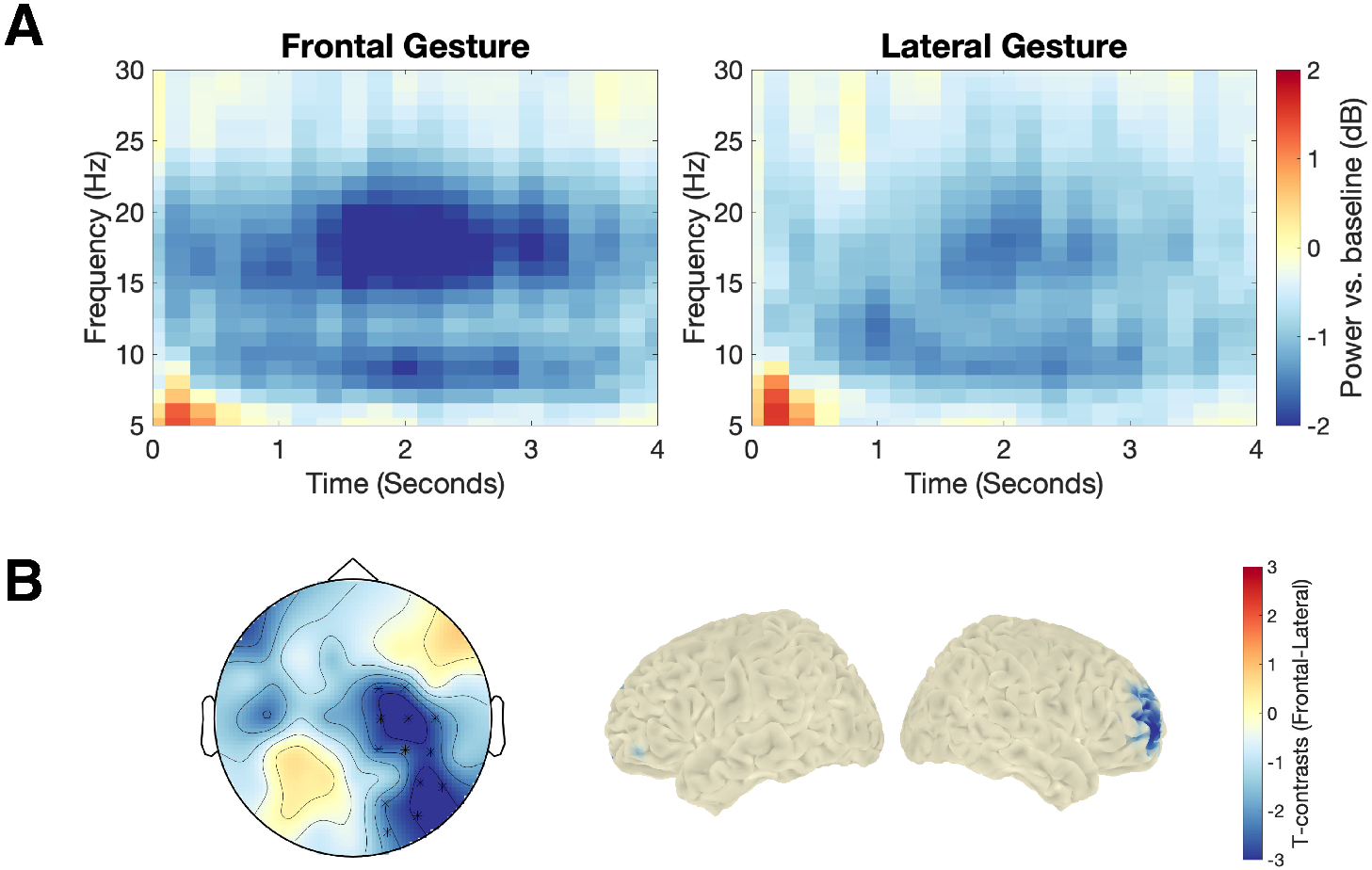
Time-frequency (TF) results at video onset for frontal and lateral gestures. Panel A: averaged TF representations across all significant electrodes showing significant difference (as in Panel B - left). Panel B (left): scalp-level t-maps for frontal vs. lateral conditions in the beta band (16-24Hz) between 0.6 to 2.4 seconds. Electrodes showing significant difference between frontal and lateral conditions (p<0.05, corrected) are marked with asterisks. Panel B (right): source-level t-maps in the beta band (16-24Hz) between 0.6 to 2.4 seconds showing significant difference (p<0.05, uncorrected) between frontal and lateral conditions.

### 3.3 Critical target word onset: N400

We compared the N400 effect firstly within each body orientation conditions (**Figure 4**). For frontal gestures, congruent vs. incongruent words elicited a significant N400 effect with a central-parietal scalp distribution (p_cluster_ = 0.006, as in **Figure 4A**). For lateral gestures, however, no N400 effect for congruency was observed (p_cluster-min_ = 0.46, **Figure 4C**). Secondly, we calculated the amplitude difference for congruency effects, and compared the congruency differences between frontal and lateral gestures to test for significance of interaction. This comparison led to a significant cluster also with central-parietal scalp distribution (p_cluster_ = 0.002).

**Figure 4.**
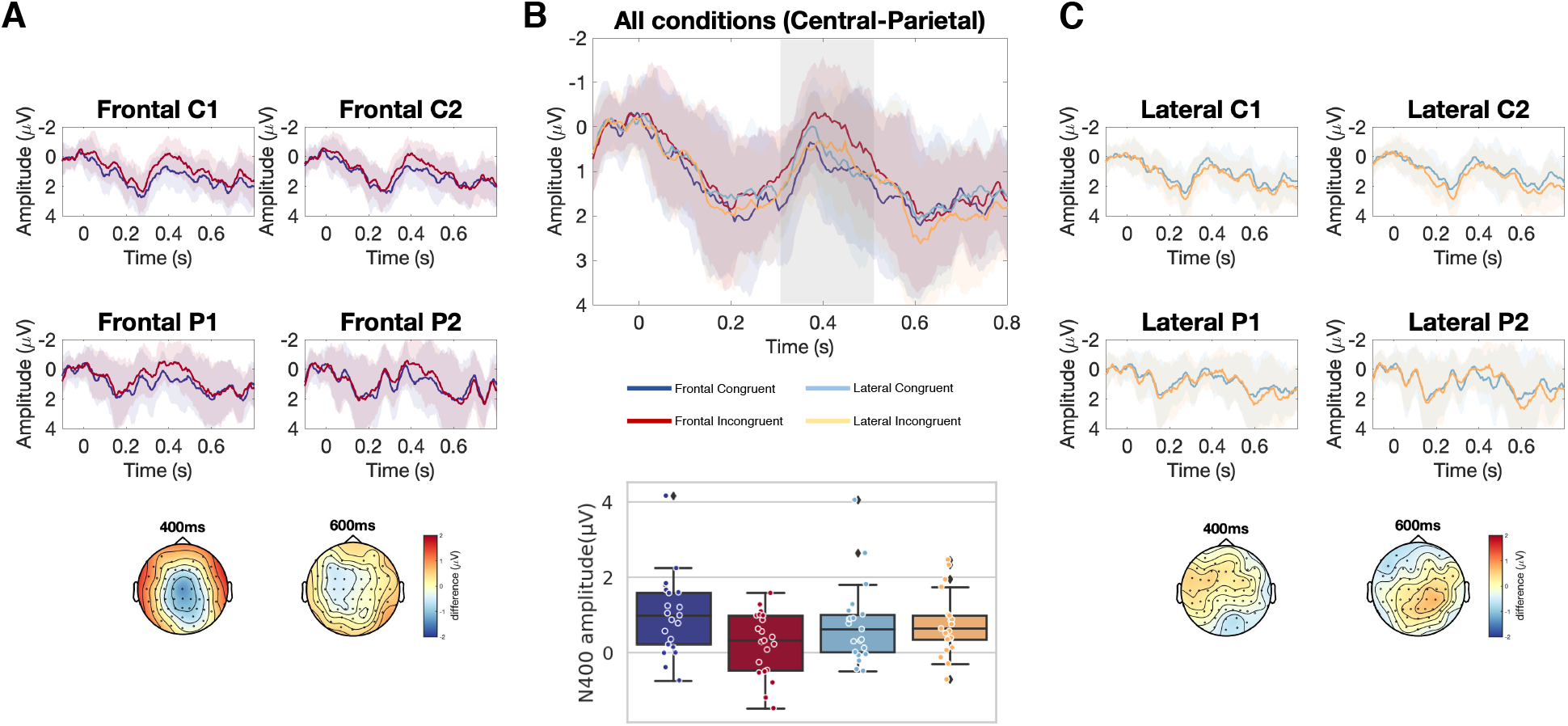
ERPs at the onset of the critical word. Panel A: ERPs for congruent vs. incongruent words primed by frontal gestures at electrodes C1, C2, P1, and P2. Scalp maps display amplitude difference at 400ms and 600ms. Panel B: ERPs averaged from nine central-parietal electrodes (C1, Cz, C2, CP1, CP1, CPz, CP2, P1, Pz, P2), and box- and swarm-plots for individual subjects’ N400 (300-500ms) amplitudes from these electrodes, for all experimental conditions. Panel C: ERPs for congruent vs. incongruent words primed by lateral gestures at electrodes C1, C2, P1, and P2. Scalp maps display amplitude difference at 400ms and 600ms. For all waveform maps, shaded areas indicate by-subject standard deviation for respective conditions.

## 4. Discussion

We conducted the current study using EEG, so as to investigate the role of body orientation on gesture observation and the semantic integration between gesture and visual sentences. During gesture observation, time-frequency analyses showed that frontal gestures elicited a more pronounced power decrease in the beta band, source localized to the mPFC. Moreover, during integration of gestures and visual sentences, we observed a clear interaction between body orientation and semantic congruency: at the critical word, the semantic congruency N400 effect observed in the frontal condition was modulated in the lateral condition. Below, we argue that these results may originate from the social-communicative nature of hand gestures.

### 4.1 Social perception of gesture’s body orientation

During gesture perception, our results showed that frontal gestures elicited a more pronounced beta band power decrease than lateral gestures. This finding not only supports the role of the beta band oscillations for perception of hand actions in general (Angelini et al., 2018; Avanzini et al., 2012; Järveläinen et al., 2004), but also corroborates the sensitivity of the beta band power to the observation of different types of gestures (He et al., 2018a; Quandt et al., 2012). Previous studies that directly tested perceptual differences of body orientation or view point of non-communicative hand actions (e.g., reaching) also showed a modulated decrease of beta power (and alpha power) for more allocentric view point (Angelini et al., 2018; Drew et al., 2015). Our study compliments this line of research by showing that this pattern could be extended to more communicative hand actions, namely gestures. However importantly, in the current study, although we observed an alpha band power decrease for both frontal and lateral gestures, unlike those studies examining non-communicative hand actions (Angelini et al., 2018; Drew et al., 2015), the alpha power did not differ between frontal and lateral orientation in our study. This data pattern attests to the hypotheses that alpha and beta power are dissociable during action or gesture observation, in the sense that alpha power decrease might be more related to sensory/motor perception in general, whereas the beta band is more sensitive to fine-grained motor-related features (Brinkman et al., 2014; He et al., 2018a; Salmelin et al., 1995).

Despite commonalities in the beta band power modulation to non-communicative hand actions, the gestures that we investigated are apparently more social. Importantly, as the beta band difference between frontal and lateral gestures were source-located to the mPFC, our finding supports the hypothesis that the difference of gesture’s body orientation is both physical and social. The mPFC has been considered as one of the most important regions for social perception and interaction (Frith & Frith, 2006), and supports the perception of social-communicative intention (Ciaramidaro et al., 2013; Schilbach et al., 2006). Previous gesture studies using fMRI also showed that the mPFC is differentially activated for frontal vs. lateral gestures (Nagels et al., 2015; Saggar et al., 2014; Straube et al., 2010), as well as gestures differing in their level of social-communicative intentions (Trujillo et al., 2019b). In line with these fMRI findings, our results provide new evidence from EEG, suggesting that frontal gestures may convey a higher degree of social-communicative intentions.

### 4.2 Social perception interacts with gesture-language semantic integration

For semantic integration between gestures and visual sentences, we observed that, at least for frontal gestures, incongruent gesture-sentence combinations elicited a classical N400 effect in terms of both latency and scalp distribution (Kutas & Hillyard, 1984). This effect concurs with an established line of literature, which suggests the functional role of the N400 during semantic integration between gesture and both verbal speech and visual sentences (Fabbri-Destro et al., 2015; Habets et al., 2011; Kelly et al., 2004; Özyürek et al., 2007; Wu & Coulson, 2005). However, in the current study, this N400 effect was modulated when participants integrate the critical word to laterally-presented gestures. This finding, to our knowledge, is a first illustration that body orientation, or potentially social-communicative aspects of gestures, may interfere with semantic integration between gesture and language. This finding supports the studies by Holle and Gunter (2007) and Obermeier et al. (2015), who observed that sentence disambiguation N400 may be modulated by 1) additional grooming gestures that are non-communicative, and by 2) watching non-grooming gestures from an actor who produces more grooming gestures. Similar behavioral findings were also reported by (Trujillo et al., 2019a), showing that gestures that are rated with differential degree of communicative intentions have a direct impact on their semantic processing.

These studies, together with our findings, suggest that higher-order social aspects of gesture may have a direct impact on the semantic processing of either visual or auditory language. In a broader sense, they imply that the established interaction between gesture and language may not only originates from their semantic, but also social-communicative nature (Holler & Levinson, 2019). Of note, the social impact of language processing was also observed by other ‘social’ manipulations, as shown by two recent studies (Jouravlev et al., 2019; Rueschemeyer et al., 2015): both studies showed that sentence processing (the N400 effect caused by contextual facilitation) could be affected by having a co-comprehender. These so-called ‘social-N400’ effects, together with the modulation effects on the N400 from our current and previous gesture studies, call for an extension of the current functional interpretation of the N400 component during sentence processing. Despite the prediction vs. integration debate (Bornkessel-Schlesewsky & Schlesewsky, 2019; Kutas & Federmeier, 2011; Lau et al., 2008; Nieuwland et al., 2019), it becomes clearer that the role of social information, especially how it interacts with semantic prediction during sentence processing, will need to be addressed by the current language processing models: Across the reported N400 studies with social manipulations, it is apparent that top-down social aspects are the only resources that has led to the reported N400 modulations, and are therefore an important component during linguistic prediction. Such an extended interactive model will not only provide a more comprehensive framework for empirical testing, but is also beneficial for a better understanding and treatment of social-communicative dysfunctions across several types of mental disorders (Dawson et al., 1998; Green et al., 2015; Yang et al., 2019).

## Acknowledgements

This research is supported by Johannes Gutenberg University Mainz. YH is supported by Johannes Gutenberg University Mainz, by the ‘Von-Behring-Röntgen-Stiftung’ (project no. 64-0001) and the DFG (project no. STR1146/11-2). B.S. is supported by the DFG (project no. STR 1146/15-1). We thank Sophie Conradi and Christiane Eichenauer for help with stimulus preparation and data acquisition.

